# *In planta* dynamics, transport-biases and endogenous functions of mobile siRNAs in *Arabidopsis*

**DOI:** 10.1101/2021.10.06.463290

**Authors:** Christopher A. Brosnan, Emanuel A. Devers, Alexis Sarazin, Peiqi Lim, Satu Lehesranta, Yrjö Helariutta, Olivier Voinnet

**Affiliations:** Department of Biology, Swiss Federal Institute of Technology (ETH), Universitätsstrasse 2, 8092 Zürich, Switzerland.; Institute of Biotechnology, University of Helsinki, PO Box 65, Helsinki, FIN-00014, Finland.; Sainsbury Laboratory, University of Cambridge, Bateman Street, Cambridge, CB2 1LR, United Kingdom.

## Abstract

- In RNA interference (RNAi), small-interfering (si)RNAs processed from double-stranded RNA guide ARGONAUTE(AGO) proteins to silence sequence-complementary RNA/DNA. Plant RNAi can propagate locally and systemically, but despite recent mechanistic advances, basic questions/hurdles remain unaddressed. For instance, RNAi is inferred to diffuse through plasmodesmata, yet how its dynamics *in planta* compares with that of established symplastic-diffusion markers remains unknown. Also unknown is why select siRNA species, or size-classes thereof, are recovered in RNAi-recipient tissues, yet only under some experimental settings. Finally, RNAi shootward movement in micro-grafted Arabidopsis – necessary to study its presumptive transgenerational effects– has not been achieved thus far and endogenous functions of mobile RNAi remain scarcely documented.
- Focusing on non-amplified RNAi in *Arabidopsis*, we show here that (*i*) transgenic RNAi-movement, although symplasmic, only partially recapitulates the diffusion pattern of free GFP *in planta*, (*ii*) the presence/absence of specific AGOs in incipient/traversed/recipient tissues likely explains the apparent siRNA-selectivity observed during vascular movement, (*iii*) stress application allows endo-siRNA translocation against the shoot-to-root phloem flow, and (*iv*) mobile endo-siRNAs generated from a single inverted-repeat(*IR*) locus, have the potential to regulate hundreds of transcripts.
- Our results close important knowledge-gaps, rationalize previously-noted inconsistencies between mobile RNAi settings, and provide a framework for functional endo-siRNA studies.

## INTRODUCTION

During RNA interference (RNAi), the RNase-III Dicer processes cohorts of 20-24-nt siRNAs from long double-stranded (ds)RNA. Loaded into AGOs, siRNAs guide endonucleolysis (‘slicing’) and/or translational repression of sequence-complementary mRNAs. Single-species of endogenous micro(mi)RNAs also are produced by Dicer from genome-encoded imperfect stem-loop precursors (Bologna and Voinnet, 2014). The present study focuses on siRNA-directed RNAi, which can also occur endogenously *via* “endo-siRNAs”. In plants, endo-siRNAs derive from *IR* loci, converging transcription or RNA-dependent-RNA-polymerase(RDR) activities (Bologna and Voinnet, 2014). The *Arabidopsis* endo-siRNA’s bulk comprises 24-nt species processed from transposable element(TE)- and repeat-derived-dsRNA by DCL3 (Xie et al., 2004), one of four Dicer-like proteins (DCL1-4). Loaded into AGO4-clade (AGO4/6/9) effectors, these guide RNA-directed-DNA-methylation(RdDM) in all cytosine contexts (CHH, CHG, CG; H: any nucleotide but G), causing transcriptional-gene-silencing (TGS) predominantly at their loci-of-origin (Law and Jacobsen, 2010). DCL4 and DCL2 produce respectively 21-nt and 22-nt endo-siRNAs from a much smaller set of *IRs and* RDR6-amplified *TRANS-ACTING-*(ta) or *PHASED-(*pha)siRNA loci(*TAS* and *PHAS*) (Gasciolli et al., 2005; Henderson et al., 2006; Liu et al., 2020). Both also convert viral dsRNA into antiviral, virus-derived (v)siRNAs (Bouché et al., 2006; Deleris et al., 2006). Like miRNAs, tasi/phasi/vsiRNAs are mainly loaded into AGO1/AGO2 to execute post-transcriptional-gene-silencing (PTGS).

In plants and some metazoans, RNAi can act non-cell-autonomously, as first recognized in tobacco displaying dynamic transgene-silencing evoking the photo-assimilates’ source-to-sink allocation path (Palauqui et al., 1996; Voinnet and Baulcombe, 1997). Graft-transmission of silencing from rootstocks to scions (Palauqui et al., 1997) was later recapitulated in micro-grafted *Arabidopsis* (Brosnan et al., 2007). Seldom examples of non-transgenic *i.e.* endo-RNAi movement include *TAS3* tasiRNAs during leaf polarization (Chitwood et al., 2009; Schwab et al., 2009), siRNAs derived from TEs or endo-*IR*s (Devers et al., 2020; Molnar et al., 2010) and phasiRNAs through inter-species grafts (Li et al., 2021). While grafting provides a simple readout of long-distance RNAi transport, its implementation and interpretations have limitations. As a major physiological stress, grafting likely exaggerates inter-organ photoassimilates’ fluxes (Kehr, 2013), which questions the significance of sRNA-target gene regulations in graft-recipient tissues (Li et al., 2021). Micro-grafting-inherent caveats also likely explain why endo-RNAi is graft-transmitted from shoot-to-root, not root-to-shoot, in *Arabidopsis*. In micro-grafts, rootstocks are made of non-photosynthetic hypocotyls and roots, whereas scions produce photosynthetic leaves. The ensuing source(scions)-to-sink(rootstocks) photoassimilates’ flux likely prevents optimal shootward RNAi-transport. The more challenging method of stem-grafting using adult plants allows shootward transport of siRNA-based phenotypes (Kundariya et al., 2020). While it imparts systemic DNA/RNA-level regulations (Lewsey et al., 2016; Molnar et al., 2010), micro-grafting-mediated shoot-to-root transmission in *Arabidopsis* conflicts with the popular notion that mobile endo-RNAi endows heritable epigenetic modifications; indeed, root apices, unlike shoot apices, do not produce a germline or embryos (Conine and Rando, 2021). Shootward movement via reproductive stem-grafting enables, by contrast, transgenerational inheritance in *Arabidopsis*, although a direct role for mobile RNAi was not strictly demonstrated (Kundariya et al., 2020). Enhancing endo-siRNA production in micro-grafted rootstocks using, *e.g.* stress, could revalorize micro-grafting for such studies by possibly enabling root-to-shoot RNAi transport against the suboptimal photoassimilates’ flux.

Heterografting between plant species/ecotypes coupled to sRNA deep-sequencing and single-nucleotide-polymorphism mapping helps discerning which, among rootstock’s and scion’s sRNAs, endow mobile RNAi (Li et al., 2021; Molnar et al., 2010). Importantly, this cannot discriminate siRNAs derived from highly conserved, repeated TEs. A common solution in *Arabidopsis* is to use homo-grafted *dcl234* triple-mutant rootstocks devoid of endo-siRNAs. Curiously, siRNA species from only some loci are retrieved in *dcl234* recipient roots (Molnar et al., 2010), suggesting that selectivity might accompany vascular RNAi movement (Maizel et al., 2020). However, the *dcl234* background itself might introduce this bias, an equally plausible, albeit untested possibility. An apparent bias also exists regarding which siRNA species, among the global products of DCL2,-3,-4 in silencing-emitting tissues, correlates with long-distance RNAi. Supporting early work on systemic transgene silencing (Hamilton et al., 2002), Molnar et al., reported that 24-nt siRNAs predominate in RNAi-recipient rootstocks of micro-grafted *Arabidopsis*. This bias was not observed, however, in a recent study of three distinct mobile RNAi sources (discussed subsequently), including via micro-grafting (Devers et al., 2020), underscoring discrepancies still unexplained to date.

In attempting to address the long-debated RNAi signal’s identity, we recently showed that AGO-free siRNAs are necessary and sufficient for cell-to-cell and long-distance RNAi movement from an *IR* transgene, an endo-*IR* locus or a naturally phloem-restricted virus (Devers et al., 2020). Moreover, deep-sequencing analyses in RNAi-incipient *versus* RNAi-recipient tissues showed that a fraction (transgene, virus) or majority (endo-*IR*) of mobile siRNAs is selectively “consumed” across traversed cells by being irreversibly loaded into cell-autonomous AGOs. A tight correlation was indeed established between siRNA cell-to-cell mobility and the 5’-nt identity of individual siRNAs, which, together with their length, largely dictates sRNA:AGO-loading specificities. Hence, 5’U/5’A-terminal siRNAs were substantially more consumed (*i.e*. less mobile) than 5’C/5’G-terminal siRNAs, respectively reflecting expression of AGO1 and AGO2/AGO4, but not of AGO5 or any known 5’G-specific AGO, in leaves (Devers et al., 2020). While its impact on vasculature-based long-distance transport remains unexplored, consumption is best accommodated in a situation where siRNAs should move via the symplasm, the cytosolic continuum shared, via plasmodesmata (PDs), by most plant cells. Indirectly supporting this notion, naturally symplasmically-isolated stomata guard-cells resist mobile RNAi (Himber et al., 2003; Voinnet et al., 1998). Moreover, mutations increasing PDs’ aperture enhance mobile transgenic RNAi (Kobayashi and Zambryski, 2007) which is decreased/increased in adult *Arabidopsis* by knockout/overexpression of receptor-like-kinases of which a subfraction is PD-associated (Rosas-Diaz et al., 2018). Nonetheless, proof that RNAi spreads symplasmically awaits the physical obstruction of PDs via spatio/temporally-controlled callose deposition, which indeed impedes endodermis-to-stele mobility of miR165a or its precursor(s) in *Arabidopsis* root tips (Vatén et al., 2011). Whether this result can be extrapolated to other miRNAs or indeed to the siRNA populations studied here remains especially unclear.

RNAi movement is often divided into distinct cell-to-cell and long-distance (*i.e.* vascular) processes, which, in reality, are likely intimately linked because the PD-based symplasmic continuum connects not only adjacent non-vascular cells but also the enucleated phloem-sieve-elements (SEs). This was elegantly confirmed in seminal studies of macromolecular transport in intact *Arabidopsis* involving freely diffusible(free-)GFP expressed from the phloem-companion-cell(CC)-specific promoter, *pSUC2* (Imlau et al., 1999; Stadler et al., 2005). Free-GFP (1) translocates via PDs from CCs-to-SEs in photosynthetically mature *i.e.* phloem-<u>So</u>urce <u>L</u>eave<u>s</u> (SoLs) in which p*SUC2* is active, (2) moves over long-distances within SEs along the transport phloem and (3) passively unloads and diffuses, via PDs, in photosynthetically immature *i.e.* phloem-<u>Si</u>nk <u>L</u>eave<u>s</u> (SiLs) in which *pSUC2* is barely active. A mobile, artificial miRNA showed distinctive differences with free-GFP in aspects of its spread (Skopelitis et al., 2018); how RNAi movement compares *in planta* with that of free-GFP remains unexplored, however.

Last but not least, the functional/biological significance of mobile endo-RNAi remains cryptic. Cell-to-cell spread of *TAS3* tasiRNAs is thought to create gene expression gradients (Chitwood et al., 2009; Schwab et al., 2009). However, this may merely reduce transcriptional repression costs over multiple cells, or regulate processes themselves non-autonomous. Likewise, while shoot-to-root transmission of endo-siRNA*-*triggered RNAi is well-established in micro-grafted *Arabidopsis,* the biological outcome of the movement of endo-siRNAs spawned from such loci is unknown. This question is particularly relevant in intact, *i.e.* non-grafted, plants given the grafting-associated caveats discussed here. In this study, we endeavoured to address most of the uncertainties remaining on mobile RNAi as laid out in this introduction in order to also close important knowledge-gaps on the process.

## RESULTS

### siRNA movement in whole plants only partially recapitulates that of free-GFP

We used *pSUC2::tmGFP9* and *pSUC2::GFP* (Stadler et al., 2005), reporting spatial patterns of respectively trans-membranous-(tm, *i.e.* cell-autonomous)GFP and free-GFP as a non-selective marker of passive macromolecular trafficking. Mobile siRNA activity was monitored side-by-side in the *pSUC2::SUL-IR* RNAi line (*SS*; Himber et al., 2003; Devers *et al*. 2020). In *SS*, PTGS of *MAGNESIUM CHELATASE*(*SUL)* causes expanded photobleaching beyond the *pSUC2* CC-specific activity domain. This reflects movement of non-amplified, AGO-free *SUL-*siRNAs (Devers et al., 2020) of which the DCL4-dependent 21-nt, species are necessary and sufficient for PTGS (Dunoyer et al., 2005). As reported, the tmGFP9 signal was CC-restricted in SoLs and below detection in SiLs where *pSUC2* activity is low; free-GFP, however, was unloaded throughout SiLs’ lamina but lowly-detectable in SoLs’ CCs (Fig.1a; Imlau et al., 1999; Stadler et al., 2005; Truernit and Sauer, 1995). The *SUL*-siRNA activity- and free-GFP movement-patterns were similar, with two exceptions. Firstly, the former was more vein-restricted in SiLs than was the latter (Fig.1a), likely reflecting *SUL*-siRNA consumption. Secondly, whereas both SoLs and SiLs displayed similarly intense vein-centred bleaching, free-GFP was strictly CC-restricted in SoLs (Fig.1a and S1). It appears, therefore, that *SUL-*RNAi spreads preponderantly into SiLs via long-distance transport and diffusion of siRNAs mainly produced in SoLs. Additionally, a mechanism apparently allows vasculature-to-mesophyll movement of siRNAs, but not of free-GFP, in SoLs.

**Figure 1.**
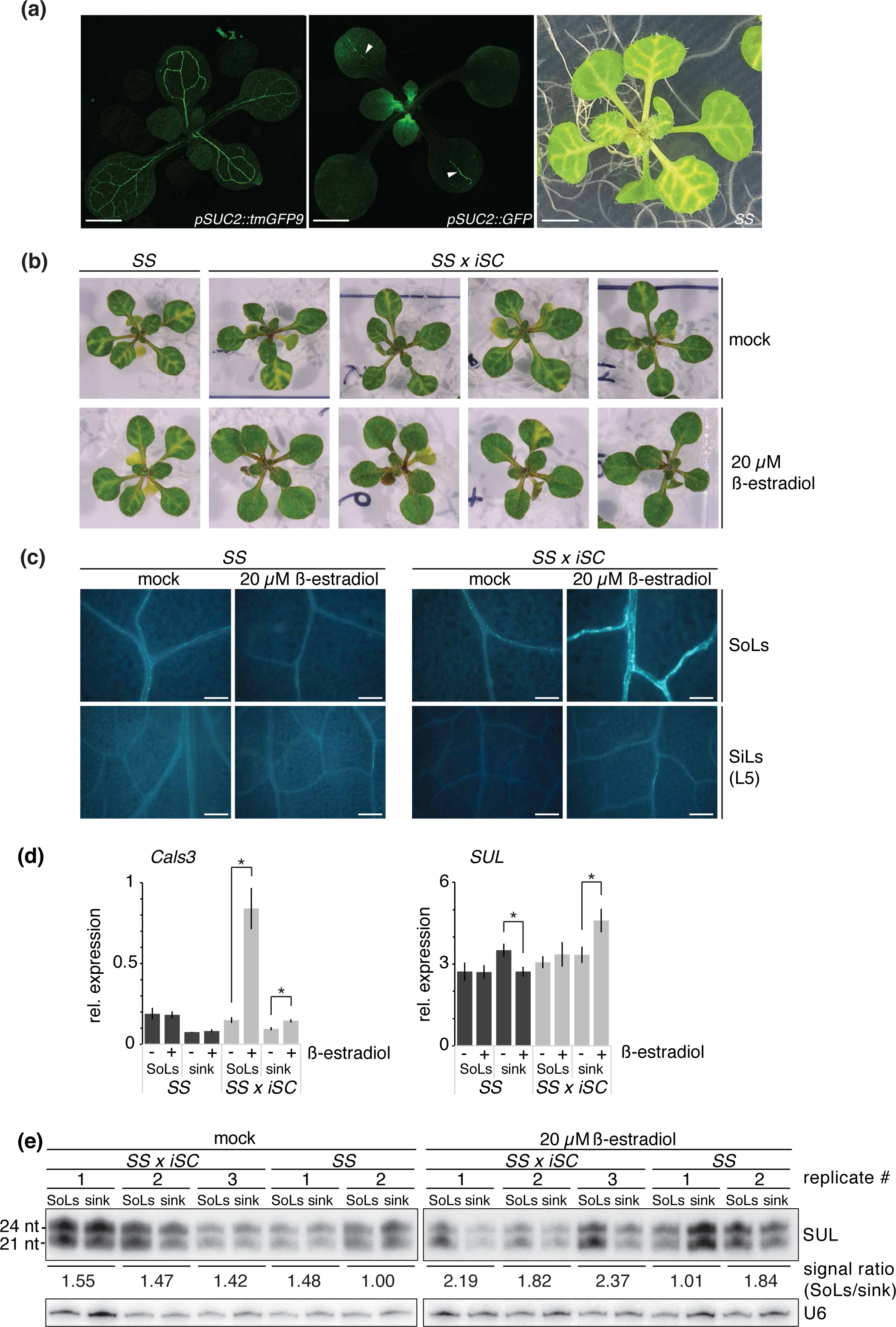
CC-to-SE translocation of the *SS*-silencing signal occurs via plasmodesmata. **(a)** Seedlings grown ½ MS containing agar plates expressing the insoluble *pSUC2::tmGFP9*, the soluble *pSUC2::GFP* or the RNAi source *SS*. Left and middle panel show fluorescence images of the respective plants. Arrowheads point to vein restriction of fluorescent signal in source leaves. Scale bar 5mm. For the phenotype of a soil grown *SS* plant, see figure 2 and S1. **(b)** 14 days old seedlings of *SS* and *SS x iSC* grown 7dpg on either DMSO (mock) or 20 µM 17-ß-estradiol containing media. See also figure S2a. **(c)** Epifluorescence microscopy analysis of callose deposition after aniline blue staining of plants taken from the experiment shown in A. Scale bars represent 200 µM. See also figure S2b. **(d)** qRT-PCR analysis of *Cals3m* and *SUL*. Black bars, *SS*; grey bars, *iSC*. Error bars represent SE, n = 6-10. Asterisks indicate statistically significant differences (p ≤ 0.05), two-tailed Student’s t-test with equal variance [F-test, p > 0.05]). **(e)** RNA gel blot analysis of source (SoLs; Cotyledons, L1-L2) and sink (L-3-7, roots) tissues from individual *SS* and *iSC* plants shown in (b). U6 serves as a loading control.

### PD occlusion in SoLs’ companion cells suffices to abrogate silencing in SiLs

The above-described *SUL*-silencing pattern would predict that targeted PD-occlusion in the SoLs’ CCs should suffice to impede bleaching in SiLs. We utilized the inducible dominant-negative iCals3m system (Vatén et al., 2011) to deposit callose at PDs’ neck regions in a CC-restricted manner. *pSUC2::XVE:iCals3m*(*iSC*)x*SS* and control *SS* seedlings were grown until full emergence of primary leaves (L1-L2), then transferred to ß-estradiol or DMSO (mock). Since mock- and iCals3m-induced *iSC*x*SS* seedlings displayed similar growth-rates (Fig. S2a), we compared RNAi movement at the L6-L7 stage. Bleaching was strongly reduced/abrogated in L3-L7 SiLs of iCals3m-induced *iSC*x*SS* plants (Fig. 1b and S2a), coinciding with substantial callose deposition in cotyledons and L1-L2 veins, but only background staining in L4-L7-veins (Fig. 1c and S2b). L3 sink-to-source transition leaves occasionally displayed partial callose deposition (Fig. S2b). These patterns agree with *pSUC2* being mostly active in SoLs (L1+L2), not SiLs (L4-L7; Fig. 1a). All mock-treated plants only displayed background staining of callose (Fig.1c). The *Cals3m* and *SUL* mRNA levels across tissues (SoLs *versus* sinks [SiLs and roots]), genotypes and treatments concurred with the aforementioned phenotypes (Fig. 1d). *DCL4* and *SUC2* mRNA levels were unchanged in either sinks or SoLs of *iSC*x*SS* plants regardless of treatments, ruling out that reduced *SUL*-siRNA accumulation underlies impaired photobleaching (Fig. S2c). While biological variation caused total *SUL-* siRNA levels to fluctuate slightly between *iSC*x*SS* plants, *SUL*-siRNA levels were consistently higher (∼40%) in source-than in sink-tissues upon iCals3m induction compared to mock treatment (Fig. 1e).

That *SUL*-siRNA retention in iCals3m-induced SoLs correlates with loss-of-silencing in SiLs supports a role for PDs at the SoLs’ CC-SE interface. It also suggests that *SUL-*siRNAs synthesized within the CCs of SoLs contributes the bulk of mobile RNAi in whole plants. Upon phloem-based long-distance transport, their passive unloading likely endows consumption-coupled PTGS in SiLs. Noteworthy, silencing was decreased in most SoLs (L1+L2) of iCals3m-induced *iSC*x*SS* plants, suggesting that the mechanism controlling siRNA-, but not free-GFP-movement outside the SoLs’ vasculature is PD-occlusion-sensitive in, at least, the CCs.

### Stem-based long-distance transport of RNAi is accompanied by minimal outflow

Both *pSUC2*-derived *SUL*-siRNAs and -artificial miRNAs (*pSUC2:amiRSUL*) were remarkably efficiently graft-transmitted into non-transgenic rootstocks (Fig.2a; (Brioudes et al., 2021; Devers et al., 2020)). Strikingly, however, photobleaching is conspicuously absent in stems in both systems (Fig. S3) and in stems of the phenotypically analogous *JAP3* line based on *pSUC2*-driven expression of a *PHYTOENE DESATURASE(PDS)-*derived *IR* transgene (Fig.2b; (Smith et al., 2007). This lack-of-silencing in stems is striking because potent *pSUC2* activity in those tissues (Stadler and Sauer, 2019) enables retrieval of apoplastic photoassimilates potentially leaked during source-to-sink transport (Ayre et al., 2003). Aiding the photoassimilates’ long-distance transport is the well-documented symplasmic isolation of the SE-CC-complex(SECCC) in stems, where few and low-conductivity PDs remaining at the CC-phloem parenchyma(PP) interface minimize photoassimilates’ dispersion from the inner vascular cylinder to outer-stem tissues (Kempers and van Bel, 1997; Kempers et al., 1998). To explore the idea that si/miRNA movement might be similarly restricted in *pSUC2*-based silencing systems, we used recombinant *Tobacco rattle virus* engineered with a *PDS* fragment (TRV-PDS), which yields photobleaching in leaves owing to virus-induced-gene-silencing (Ratcliff et al., 2008). Because viral movement proteins (MPs) actively enlarge PDs’ aperture including at the stems’ CC-PP-interface (Itaya et al., 2002), we anticipated that TRV-PDS-infected stems would display extensive photobleaching, which was indeed the case (Fig. 2b). These results support the notion that the SE-CC’s isolation efficiently limits si/miRNA outflow and, hence, dilution, during their stem-based long-distance transport. This possibly explains the graft-transmission’s efficacy and potent systemic effects of *SUL*-siRNAs despite their mostly SoL-based production (Fig. 2a).

**Figure 2.**
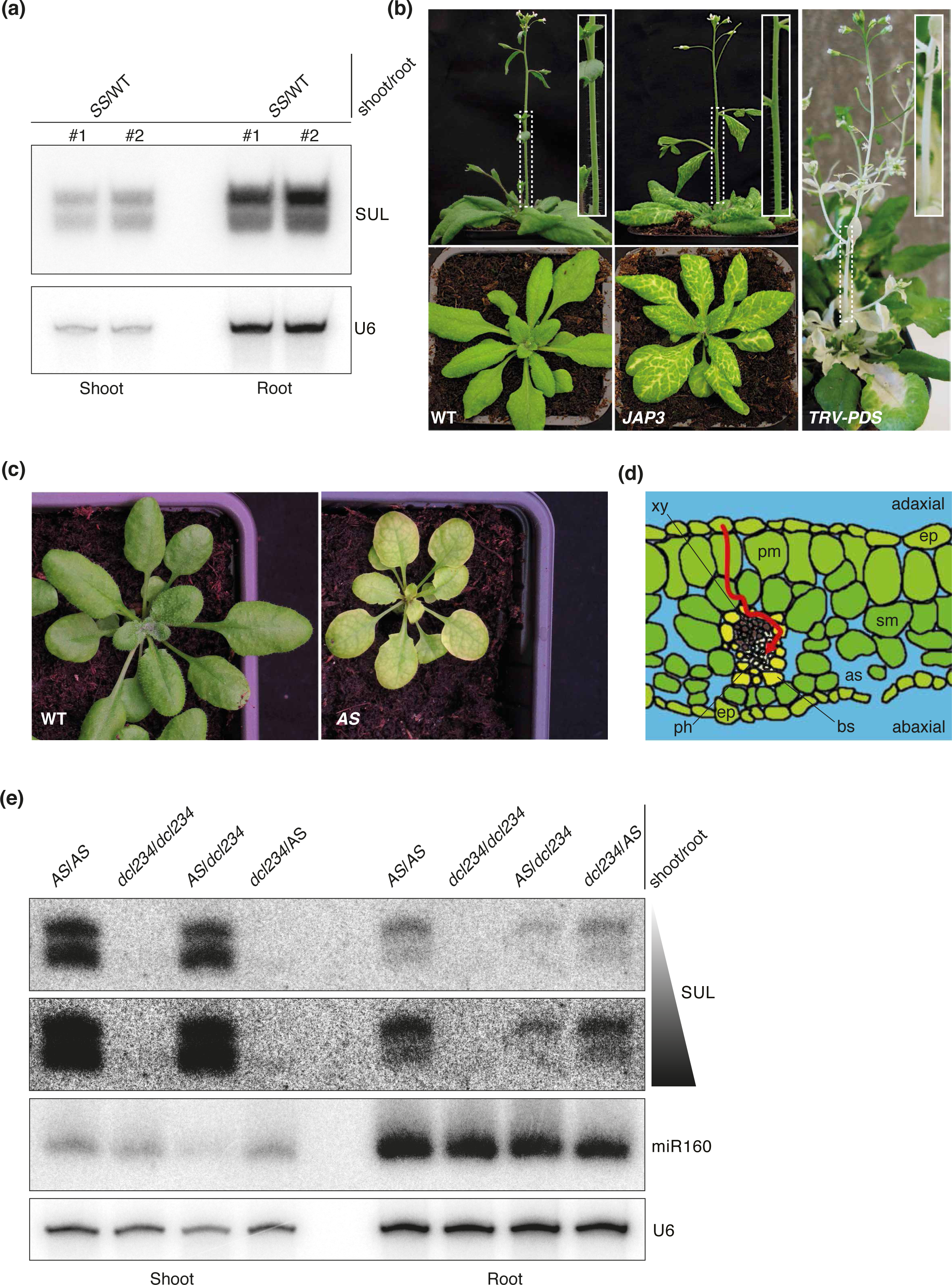
Consumption, initiating expression domaining and PD architecture dictate the mobile nature siRNA species. **(a)** RNA gel blot analysis of biological duplicates of *SS* shoot grafted to WT root. U6 serves as a loading control. **(b)** Images of equivalent age rosette stage (bottom panels) and flowering (top panels) WT (left) and *JAP3* (middle panels) plants. Right panel shows TRV-PDS infected flowering plant. Insets show distinctive lack or presence of bleaching in stem tissues. **(c)** Images of whole plant showing photobleaching phenotypes of WT and *AS* lines. **(d)** Cross section of an *Arabidopsis* leaf showing the distance siRNAs would have to travel from adaxial or abaxial sides to initiate vasculature-based graft transmissible silencing (red arrow). The expression domain of ATML1 is exclusively located in the epidermis. Abbreviations: as, air space; bs, bundle sheath; ep, epidermis; ph, phloem; pm, palisade mesophyll; sm, spongy mesophyll; xy, xylem. **(e)** RNA gel blot analysis of shoots (left; lane 1-4) and root (right; lane 6-9) of indicated graft combinations. Black triangle indicates increase in exposure for the SUL probe. miR160 and U6 serve as loading controls.

### Differential AGO-mediated consumption across vein-distal incipient tissues likely causes unequal siRNA-length representations in micro-grafted recipient tissues

Both the 21-nt and 24-nt *SUL*-siRNA species were graft-transmitted into rootstocks (Fig. 2a), contrasting with a previously-established bias towards 24-nt siRNA long-distance mobility (Molnar et al., 2010). To test if differential AGO-mediated consumption between experimental systems could explain this discrepancy, we engineered *pATML1::SUL*(*AS*) *Arabidopsis*, in which the epidermis-specific promote*r ATML1* drives the *SS*-*IR*. *AS* leaves accumulate equal amounts of 21-nt and 24-nt *SUL*-siRNAs but display laminal surface-bleaching likely reflecting *SUL*-siRNA movement from the epidermis to the lower, photosynthetically-active mesophyll (Fig. 2c). In contrast to *SS*, where CC-produced siRNAs are directly delivered into SEs, more consumption would be expected for *AS*-derived siRNAs during their multi-cell-layer epidermis-to-SE movement via the mesophyll (Fig. 2c-d). Indeed the vasculature remains green in *AS* leaves (Fig. 2c). Accordingly, and unlike the relatively unbiased transmission of *SS-*derived 21- and 24-nt *SUL*-siRNAs (Fig.2a; Devers et al., 2020), *AS-*derived 24-nt *SUL*-siRNAs predominated in micro-grafted rootstocks (Fig. 2e), evoking the previously-reported 24-nt siRNA bias (Molnar et al., 2010). This suggested a strong and selective consumption of the *AS-*derived 21-nt *SUL*-siRNAs in silencing-emitting scions. Accordingly, the transcript of *AGO4* (loading the 24-nt *SUL*-siRNAs; Devers et al., 2020) is ∼3 and ∼7 times less abundant that of *AGO1* (loading the 24-nt *SUL*-siRNAs Devers et al., 2020) in *Arabidopsis* whole leaves and mesophyll, respectively (Fig. S4). Thus, the 24-nt siRNA bias in the *AS* system most likely reflects high and selective consumption of 21-nt siRNAs rather than selectivity in long-distance movement *per se*. The results also support the expected, albeit so-far-untested notions that (i) sRNA size, in addition to 5’-nucleotide identity, strongly influences AGO-mediated consumption, and that (ii) consumption affects not only cell-to-cell but also long-distance vascular movement.

### *dcl234* rootstocks are suboptimal RNAi-recipient tissues in micro-grafted *Arabidopsis*

By micro-grafting *SS* scions onto siRNA-deficient *dcl234* rootstocks, we could directly compare the shoot-to-root spread of *SUL-*siRNAs with that of endo-siRNAs. These included the DCL2/DCL3-dependent 22-nt/24-nt siRNAs from the *IR71* single-locus (Henderson et al., 2006) and two TE/repeat-derived-species (*rep2*,*siRNA1003;* Fig. 3a). All siRNAs moved into micro-grafted *dcl234* rootstocks, albeit to varying extents (Fig. 3a-b;(Devers et al., 2020)). *SUL*-siRNA-was stronger than *IR71*-siRNA graft-transmission, likely reflecting higher production levels of the former. Nonetheless, the 22-nt and 24-nt *IR71*-siRNAs accumulated equally in *dcl234* rootstocks, suggesting limited consumption. This was consistent with *IR71*, like *SS*, but unlike *AS*, being expressed within the leaves’ vasculature of RNAi-emitting scions (Devers et al., 2020). Both *rep2* and *siRNA1003* siRNAs accumulated efficiently in *dcl234* rootstocks, contrasting with previous deep-sequencing showing a very low (0-33%) shoot-to-root transmission of these and additional 24-nt endo-siRNAs (Molnar et al., 2010, reanalysed in Fig. S5). Sequencing biases aside, the phloem flow was perhaps better restored under our micro-grafting conditions. Alternatively, AGO4/AGO6 destabilization due to lack-of-cargoes in siRNA-deficient *dcl234* recipient rootstocks – as originally reported in *rdr2* and *nrpd1a* (Havecker et al., 2010; Li et al., 2006) – was possibly less pronounced as was, perhaps, its reciprocal consequence, *i.e.* the enhanced turnover of non-loaded 24-nt siRNAs (Havecker et al., 2010; Smibert et al., 2013; Vaucheret et al., 2006). We considered these possibilities noteworthy because AGO-loaded species contributes mainly to siRNA deep-sequencing results (Grentzinger et al., 2020).

**Figure 3.**
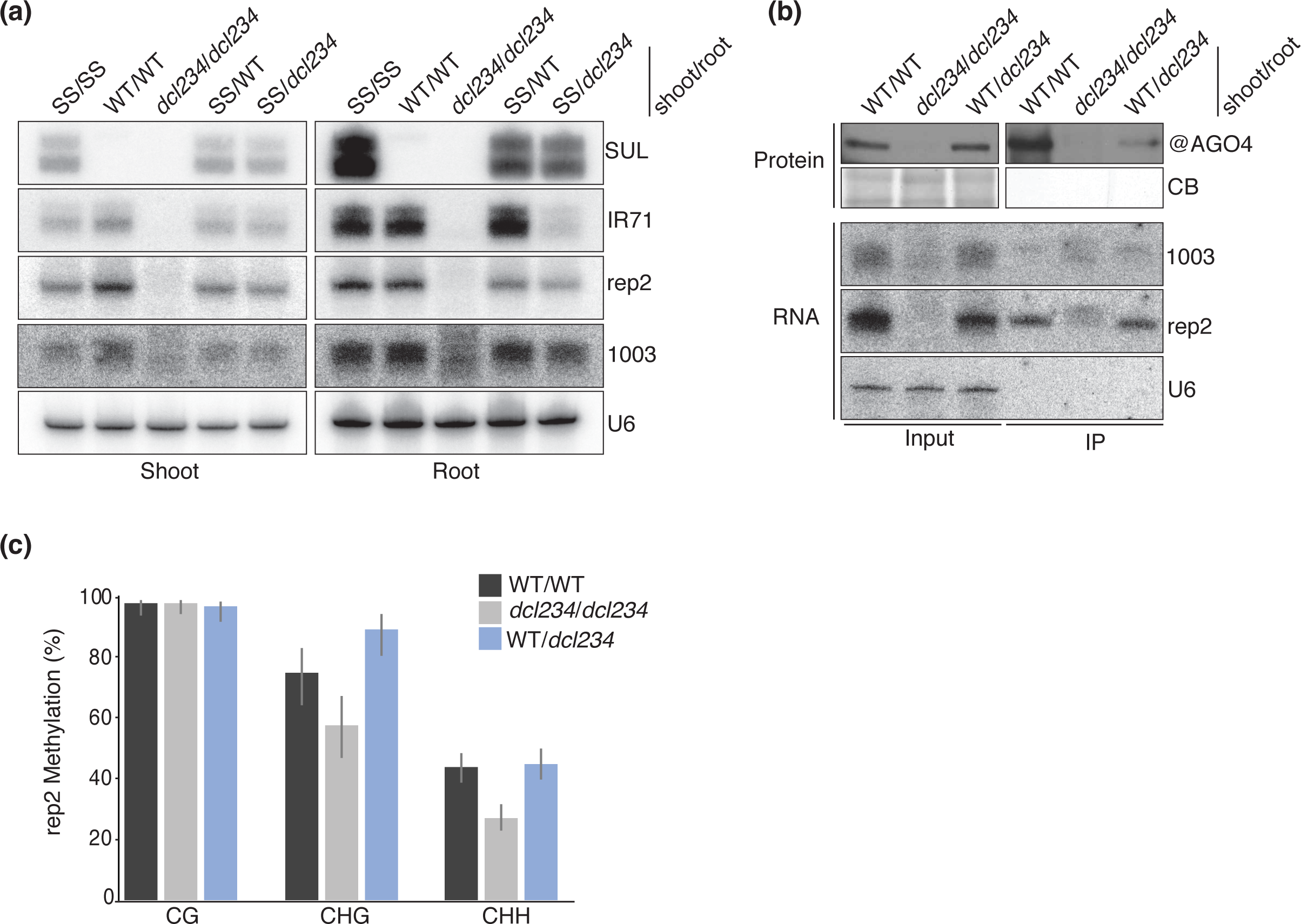
Graft-transmissible endo-siRNA movement, limiting factors and functionality. **(a)** RNA gel blot analysis of the indicated graft combinations looking at either transgene- (*SS*) or endo-siRNA movement to a WT or *dcl234* root. U6 serves as a loading control **(b)** Immunoprecipitation of AGO4 in the indicated graft combinations. Top panels represent western blot analysis of AGO4 protein levels Input (Left) and IP (Right). Coomassie Blue (CB) serves as a loading control. Lower panels represent RNA gel blot analysis of the same immunoprecipitations. U6 serves as a loading control **(c)** Bisulfite sequencing of *rep2* loci in the of the root of the indicated graft combinations. Graphs depict averages with error bars showing 95% Wilson score confidence intervals. Numbers and distribution are shown in Table S1.

Consistent with AGO4 reduced accumulation impairing siRNA detection in micro-grafted *dcl234* recipient tissues, AGO4 levels were partially rescued in rootstocks of WT/*dcl234 versus dcl234*/*dcl234* micro-grafted plants (Fig. 3b). Thus, upon translocation into *dcl234* rootstocks, shoot-derived 24-nt siRNAs had likely partially stabilized AGO4 via loading. Accordingly, *rep2* and *1003* siRNAs were readily detected in AGO4 IPs performed in rootstocks of WT/*dcl234* unlike *dcl234*/*dcl234* plants (Fig. 3b). The AGO4-destabilization’s impact of *dcl234* was revealed by activity-measurements of graft-transmitted *rep2* siRNAs (Fig. 3c). Methylation at *rep2* was reduced from 75.64%-to-58.33% and from 44.23%-to-27.55% at CHG and CHH sites, respectively, in WT/WT *versus dcl234*/*dcl234* grafted rootstocks (Fig. 3c). siRNAs from WT shoots grafted onto *dcl234* rootstocks (WT/*dcl234*) rescued CHG- and CHH-methylation (Fig. 3c). As expected, CG-methylation remained unchanged, consistent with its siRNA-independent maintenance (Fig. 3c; Xie et al., 2004). Coincident AGO4 re-stabilization and DNA methylation-rescue confirm functional movement of 24-nt endo-siRNAs, yet at levels and with activities likely underestimated by use of suboptimal and bias-inducing *dcl234* recipient tissues owing to an artefactual paucity of AGO4-clade RdDM-effectors. Without altering their graft-transmission *per se*, this caveat probably decreases detectability of low-to-moderately abundant mobile 24-nt siRNAs.

### Stress-induced shootward movement of endo-siRNAs in micro-grafted *Arabidopsis*

In reverse-grafting, *SUL-*siRNAs moved from *SS* transgenic rootstocks to WT shoots, albeit less efficiently than from shoot-to-root (Fig. 3a and 4a). Deep-sequencing confirmed this observation (Devers et al., 2020; Fig. S6a) reminiscent of similar transmission-deficits of transgene-derived and endo-siRNAs under similar root-to-shoot micro-grafting settings (Molnar *et al*. in 2010). To explore a comparable endo-siRNA transmission-deficit without the AGO4-destabilization effect of *dcl234*, we exploited an *IR71* T-DNA knockout (*ir71*) essentially eliminating *IR71*-siRNAs (Fig. 4b and S6b); these are near-identical to those of *SS* in their vasculature-proximal production, mobile attributes and RDR-independency (Devers et al., 2020). Like *SUL*-siRNAs, *IR71*-siRNAs were at northern detection limit in shoots of *ir71*/WT grafts (Fig. 4c, control) likely due to suboptimal root-to-shoot photoassimilates’ fluxes incurred by micro-grafting. We thus tested if this caveat could be overcome by increasing *IR71*-siRNA production through stress application. Among several stresses tested, heat-shock (20 hrs at 37°C; Fig. 4d) caused a ∼4-fold increase in *IR71* promoter activity (Fig. 4e). In side-by-side experiments, HS applied under otherwise identical conditions to those of mock-treated plants caused detectable accumulation of 22-nt and 24-nt *IR71*-siRNA in recipient *ir71*-shoots (Fig. 4c, 37°C). Therefore, by stimulating *IR71*-siRNA production/mobility, HS could overcome a major micro-grafting limitation.

**Figure 4.**
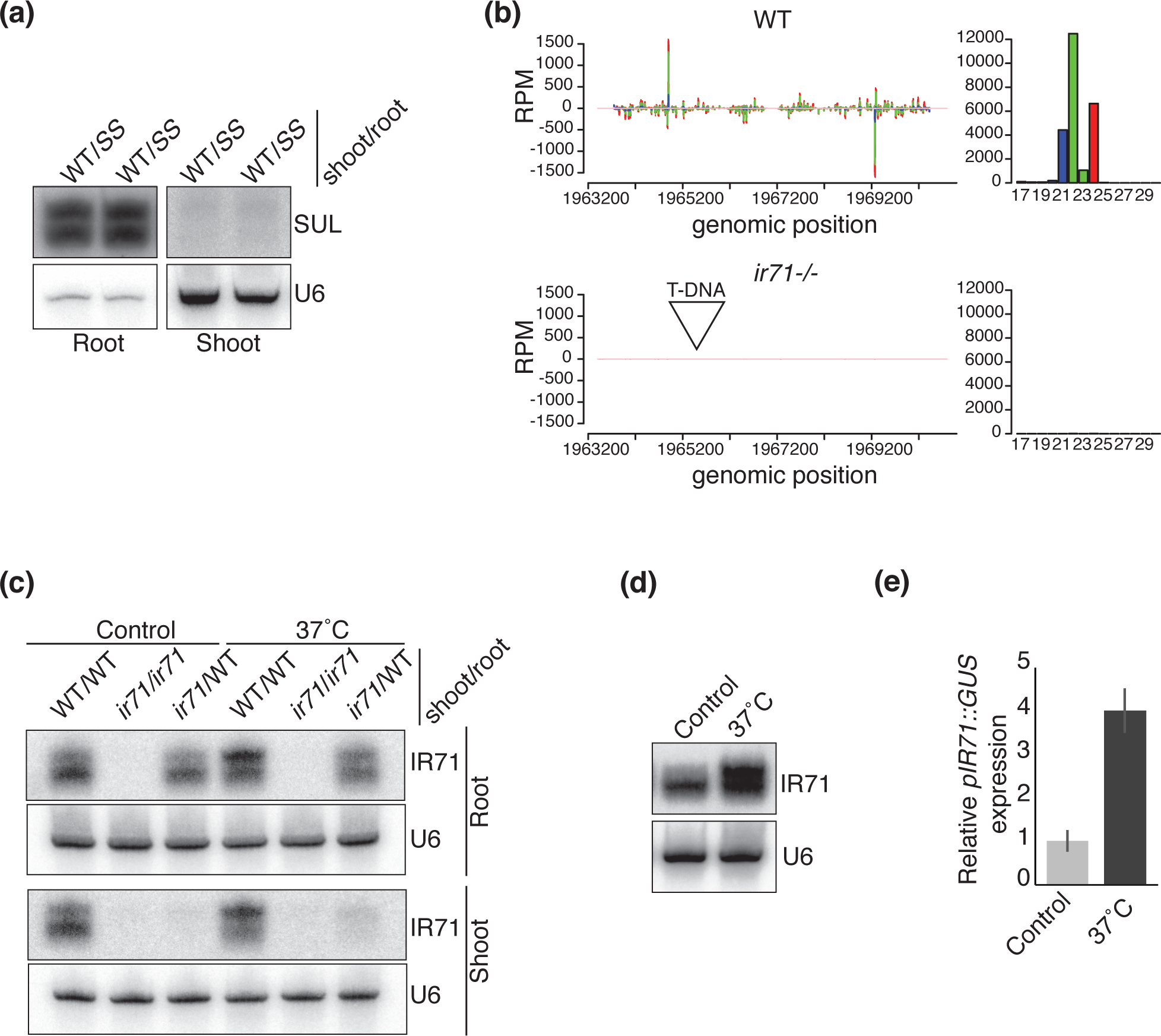
Heat shock inducible root-to-shoot movement of *IR71* derived endo-siRNAs. **(a)** RNA gel blot analysis of Root (left) and Shoot (right) of SS root grafted to WT shoots. U6 serves as a loading control. **(b)** siRNA profiles of IR71 loci in WT and ir71 mutant lines (see also figure S5b). Upper panels show the profile and size distribution (right) of IR71-derived siRNAs in WT plants. Lower panel shows siRNA profile of IR71-derived siRNAs in T-DNA line (*ir71 -/-*). RPM – Reads per Million, colour code: Blue – 20-21-nt, Green – 22-23-nt and Red – 24-25-nt. **(c)** RNA gel blot analysis of inflorescence tissue under heat shock (HS) conditions for the indicated time. U6 serves as a loading control. **(d)** qRT-PCR analysis of heat shocked plants expressing a *pIR71::GUS* reporter. Error bars represent +/- SD of 3 biological replicates. **(e)** RNA gel blot analysis of indicated graft combinations from either the root or scion of *IR71*-derived siRNAs under control or heat-shock conditions.

### Mobile *IR71*-siRNAs mediate *cis*-methylation and negatively regulate dozens of transcripts linked to systemic acquired resistance in intact plants

As much as the physiological relevance of grafting to (endo)-siRNA mobility is debated (Kehr, 2013; Liang et al., 2012) so too is the biological role, if any, of non-cell autonomous silencing mediated by these molecules in intact (*i.e.* non-grafted) plants, aside from *TAS3* siRNA cell-to-cell movement during leaf polarization (Chitwood et al., 2009; Schwab et al., 2009). To address this key issue we exploited our previous findings that (i) *IR71* is vascularly-expressed and (ii) mobile *IR71*-siRNAs can be physically isolated, from the lower epidermis located several cell-layers away ((Devers et al., 2020); Fig. 5a). We demonstrated on multiple occasions the tissue-specific mechanical separation including in the case of *IR71* (Brioudes et al., 2021; Brosnan et al., 2019; Devers et al., 2020). Bisulfite sequencing of *IR71* DNA from the incipient vasculature revealed high levels of CG (80%) and moderate levels of CHG and CHH methylation (∼50% and ∼10% respectively; Fig. 5b). By contrast, methylation was markedly reduced at asymmetric, RdDM-dependent cytosines in both vasculature and distal epidermis of *ir71* mutant leaves (Fig. 5b). Furthermore, HS treatments slightly increased *IR71*-siRNA movement and asymmetric cytosine methylation in the epidermis (Fig.5a and c). Therefore, mobile *IR71*-siRNAs mediate RdDM *in cis* in intact plants.

**Figure 5.**
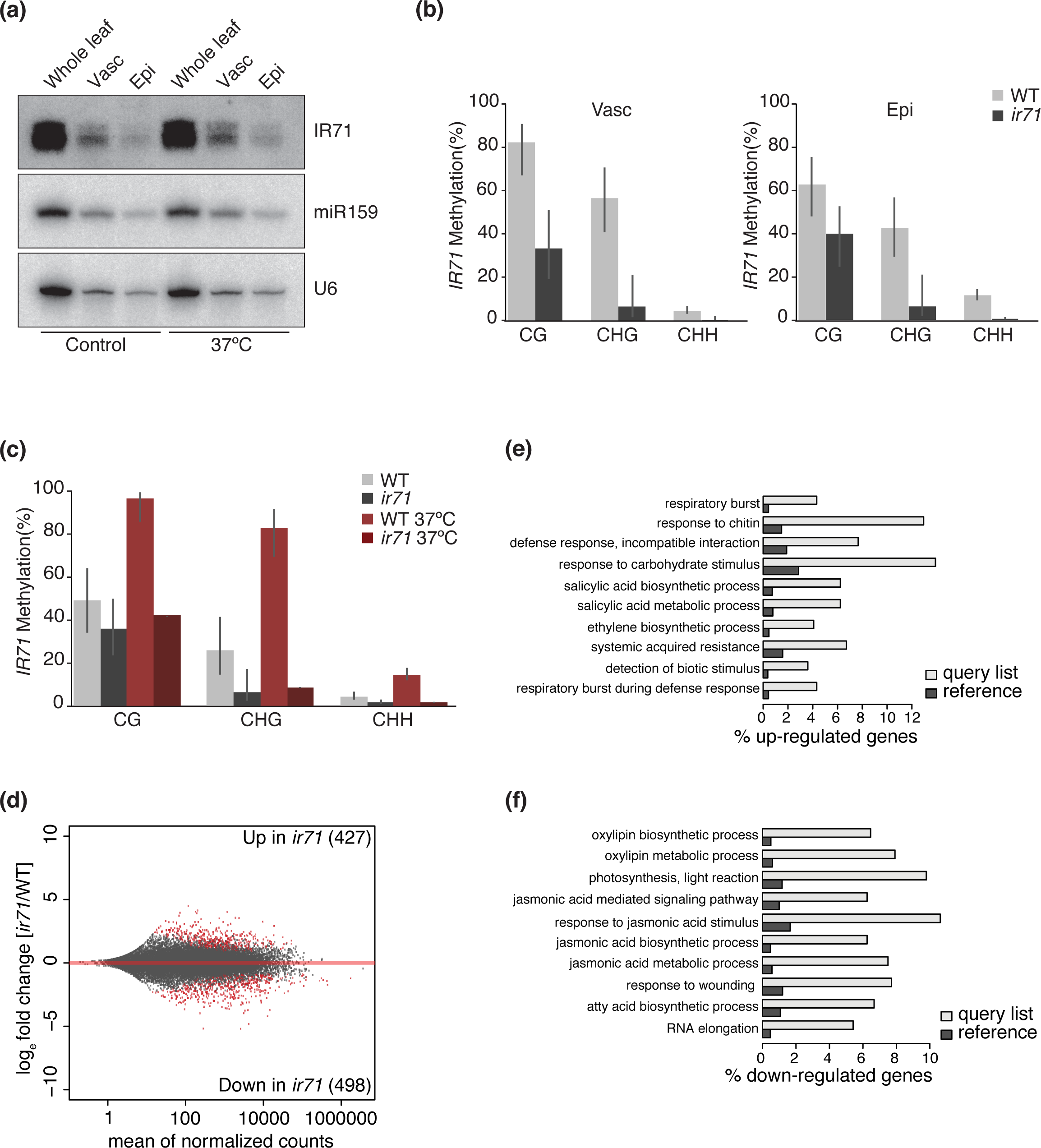
Functionality of endogenous inverted-repeat derived siRNAs under native conditions. **(a)** RNA gel blot analysis of *IR71*-derived siRNAs from the either whole leaf, vasculature (Vasc) or epidermis (Epi) under control (left) or heat shock (right; 37°C). miR159 and U6 serve as controls. **(b)** Bisulfite sequencing analysis of *IR71* locus from the vascular (Vasc) and epidermal (Epi) tissue in either WT or *ir71* mutants. Graphs depict averages with error bars showing 95% Wilson score confidence intervals. Numbers and distribution are shown in Table S1. **(c)** Bisulfite sequencing analysis of *IR71* locus from epidermal tissue either Wildtype (Col) or *ir71* mutants under normal or heat shock (37°C) conditions. Graphs depict averages with error bars showing 95% Wilson score confidence intervals. Numbers and distribution are shown in Table S2. **(d)** MA-plot representing the result of mRNA-seq differential analysis between epidermal tissue of WT or *ir71* mutant. The number of genes considered as significantly up- or down-regulated (red dots; FDR < 0.01) are shown on the top- or bottom-right. **(e)** Top ten GO-term enrichments of the 427 upregulated genes. **(f)** Top ten GO-term enrichments of the 498 downregulated genes.

To explore the global *trans*-regulatory impact of *IR71*-siRNAs – encompassing both PTGS and TGS levels – under native (*i.e.* non-grafted) conditions, we conducted whole-transcriptome analyses of Meselect-isolated epidermis of *ir71 vs* WT leaves to focus solely on the locus’ non-autonomous regulatory effects. In high-stringency analyses (FDR cut-off ≤ 0.01) 427 and 498 transcripts were respectively up- and down-regulated in the *ir71 vs* WT epidermis (Fig. 5d). GO-term-enrichment of up-regulated mRNAs identified processes strikingly dominated by pathogens’ recognition and salicylic acid (SA) biosynthesis/regulation/signalling (Fig. 5e). SA production is triggered during plant-biotroph incompatible interactions following a respiratory burst culminating in the “hypersensitive response” (HR). SA accrues not only within and around HR lesions, but also in distant, non-infected tissues where resistance to secondary infection is primed via “systemic acquired resistance” (SAR; Fu and Dong, 2013). By contrast, processes of biosynthesis/signalling by the oxylipin jasmonic acid (JA) –conferring resistance to necrotrophs and herbivores/wounding– prevailed in the down-regulated mRNA set (Fig. 5f). This agrees with SA *vs* JA signalling being reciprocally antagonistic (Pieterse et al., 2009) and provides the first comprehensive view of gene expression changes caused solely by mobility of a genetically-tractable endo-siRNA cohort in intact plants. Given its large, yet specific mobile regulatory output, *IR71* therefore displays attributes of a non-cell autonomous modulator of SAR.

## DISCUSSION

### Systemic RNAi in *pSUC2*-based systems is the manifestation of mostly a single movement process

We conclude from our study that mobile RNAi in young tissues of *SS* and related *pSUC2*-based systems (Smith et al., 2007; Uddin et al., 2014) reflects diffusion of an siRNA-pool produced within, and translocated from, mature tissues, rather than active and localized cell-to-cell movement in young tissues. Callose deposition in SoLs’ CCs indeed suffices to abrogate RNAi-movement in *SS* plants (Fig. 1). As highlighted by (Paterlini et al., 2021), high callose accumulation within iCals3m-expressing CCs could, in principle, also interfere with apoplastic transport. Under the iCals3m-setting, however, sucrose/trehalose-6-phosphate transport was unaffected (Paterlini et al., 2021), and, likewise, plant growth-rates did not differ here between treatments/genotypes (Fig. S2a). Therefore, PDs are likely central to source-to-sink RNAi translocation, primarily at the SoLs’ CC-SE interface. This rationalizes why Cadmium treatment in SoLs inhibited long-distance but not cell-to-cell transgene silencing (Ueki and Citovsky, 2002) since heavy metals induce callose deposition specifically in vasculature-associated cells (Ueki and Citovsky, 2005). Upon phloem unloading, the large PD size-exclusion-limit in SiLs (Stadler et al., 2005) would accommodate passive diffusion of translocated ∼13-15 kDa siRNAs (21-nt duplexes) well-below the free-GFP’s 27 kDa. RNAi movement, at least in *pSU2*-based systems, is thus the likely manifestation of predominantly vasculature-dependent long-distance siRNA transport, as indeed previously anticipated (Stadler and Sauer, 2019).

### Differential AGO-mediated consumption of siRNAs in distinct experimental settings likely explains an apparent size-selectivity in their mobility

Free-GFP invades the near-entirety of SiLs’ laminas, reflecting that no conceivable plant-based mechanism might hinder this xenobiotic protein’s spread. AGO-siRNA co-evolution, by contrast, likely underpins the vein-restricted *SUL*-siRNA activity-pattern in SiLs. Of the 21-nt and 24-nt species only the former mediates mobile PTGS/photobleaching in an AGO1-(5’U) and, secondarily, AGO2-(5’A)-dependent manner (Devers et al., 2020). Of the two, AGO1-based consumption likely prevails due to its overrepresentation in leaves (Fig. S4). Accordingly, genetically reducing AGO1 loading-efficacy in silencing-traversed cells suffices to extend 5’U-si/miRNA physical movement throughout leaves (Brioudes et al., 2021; Devers et al., 2020). Loaded into the lesser abundant AGO4, a higher proportion of 24-nt *SUL*-siRNAs would likely move beyond the vein-restricted activity-pattern of their 21-nt counterparts. This would be undiagnosed, however, as they do not contribute PTGS/photobleaching (Dunoyer et al., 2005). In the *AS* setting, multiple cell-layers separating incipient epidermis from recipient SEs likely exacerbate the differential AGO1 *versus* AGO4 consumption, rationalizing how 24-nt *SUL*-siRNAs are near-exclusively detected in graft-recipient tissues (Fig. 2e). Consistent with the above observation, an artificial *ATML1-*driven 5’U-terminal miRNA displayed no apparent vascular activity unless expressed at extremely high levels presumably overcoming its AGO1-mediated consumption across epidermis-to-vasculature-traversed tissues (Brioudes et al., 2021; Skopelitis et al., 2018).

Consumption also resolves the long-standing conundrum as to why 24-nt siRNAs appear selectively correlated with long-distance mobility in some, unlike other, experimental systems. In GFP-transgenic *N.benthamiana*, *Agrobacterium*-based transient expression of silencing-inducing constructs initiate mobile RNAi. Both 21-nt and 24-nt *GFP*-siRNAs accumulate mostly in the agroinfiltrated patch’s epidermis and mesophyll (Hamilton et al., 2002) where high *AGO1* levels (Fig. S4) and thus strong AGO1-based consumption likely prevents 21-nt *GFP*-siRNAs from reaching the vasculature. This likely explains why 24-nt *GFP*-siRNAs, presumably far-less AGO4-consumed, positively correlate with systemic silencing in this system (Hamilton et al., 2002). The strong 24-nt siRNA bias observed genome-wide in the micro-grafting studies of Molnar *et al*. (2010) presumably also reflects that only a subfraction of the total pool of 21-nt endo-siRNA species is sufficiently vasculature-proximal to evade significant consumption by AGO1. Indeed, an siRNA-length bias was not observed in three-out-of-three independent systems studied in Devers *et al*. (2020), all of which were artificially (*SS*) or naturally (polerovirus and *IR71*) poised to trigger mobile RNAi in the direct SEs’ vicinity, thereby mostly bypassing AGO1-mediated consumption. Thus, what might be interpreted as selectivity/conditionality based on mobile siRNA-size most likely reflects system-intrinsic differences influencing the extent of differential AGO-mediated consumption. From a natural standpoint, these findings reinforce the previously-made suggestion (Skopelitis et al., 2018) that efficient long-distance movement would be an attribute of si/miRNAs naturally expressed close to, or within, the vasculature, as are indeed all systemic miRNAs reported so far in plants (Brioudes et al., 2021; Brosnan et al., 2019; Lin et al., 2008; Pant et al., 2008).

### High efficiency of long-distance RNAi transport

It has been proposed that vascular miRNAs might only display efficient non-cell autonomous effects if their activity is fostered, in recipient tissues, by “transitivity”, whereby mobile secondary siRNAs are processed from long dsRNA amplified by RNA-DEPENDENT-RNA-POLYMERASES(RDRs) from cleaved miRNA-target transcripts (Skopelitis et al., 2018). However, long-distance movement of miR399 (Lin et al., 2008; Pant et al., 2008), miR2111 in *Lotus japonicus* (Tsikou et al., 2018) and amiRSUL (Brioudes et al., 2021) is not accompanied by secondary siRNA production. We proposed, similarly, that primary *i.e.* non-RDR-amplified RNAi might suffice for robust long-distance siRNA-mediated silencing (Devers et al., 2020), even though it had been so far impossible to disrupt, in real time, ongoing RNAi *in planta*, a necessary approach to measure the process’ potency and persistence. Our ability to interrupt *SS-*silencing via targeted callose deposition now reveals that transport initiated in only 2-3 SoLs is sustained, without RDR amplification, in at least four SiLs. Additionally, none-of-three *pSUC2*-based si/miRNA silencing systems displayed stem photobleaching in contrast to the viral-based TRV-PDS-infected stems. This suggests that symplasmic isolation of stem SE-CC complexes –which is disrupted by virus-encoded MPs– ensures optimal long-distance transmission of mobile si/miRNAs by limiting/precluding their dilution in stems. This efficacy, in turn, rationalizes why transitivity is dispensable in recipient sink tissues, in at least the aforementioned mi/siRNA systems and during cross-kindom (Subhankar et al., 2021). We note, however, that *SUL* silencing in SiLs likely requires a continued RNAi input because residual *SUL-*siRNAs in SiLs failed to promote bleaching after PD closure in SoLs (Fig. 1). Thus, transitivity might primarily enable silencing sustainability over prolonged periods of time. This might be particularly relevant during (often life-long) viral infections (Pumplin and Voinnet, 2013) or extended developmental processes such as leaf polarization (Chitwood et al., 2009; Schwab et al., 2009). By contrast, transitivity may not be a desirable feature of short-term sRNA-based responses requiring rapid reversal, for instance upon the elapse of a stress.

### Possible active forms of cell-to-cell RNAi transport

PD closure at the CC-SE interface reduced bleaching not only in the signal-recipient SiLs, but also in the signal-incipient SoLs in which, therefore, *SUL-*siRNAs must also move via PDs toward the mesophyll at the CC-PP-bundle-sheath interface. Similar movement is observed in *pSUC2*-based artificial miRNA systems, including *amiRSUL*, though the involvement of PDs has not been experimentally verified in these cases (Brioudes et al., 2021; Skopelitis et al., 2018). In sharp contrast, free GFP remains strictly confined within the CCs in SoLs (Fig. 1a and S1; (Brioudes et al., 2021; Skopelitis et al., 2018), in line with its retention in phloem cells where only apoplastic transport occurs (Stadler et al., 2005; Werner et al., 2011). Thus, a mechanism exists in SoLs, that propagates si/miRNAs outside the CCs against the photoassimilates’ flux in a manner not permitting free GFP movement. Moreover, this process must occur, at least partly, at the CC-PP interface in which low-conductivity PDs occur at a low frequency, incidentally explaining the prevailing apoplastic phloem-loading of *Arabidopsis* (Turgeon, 1996). Therefore, the CC-to-PP sRNA movement in SoLs is perhaps the manifestation of an active PD-based process, unlike movement seen at most other cellular junctions involved in *pSUC2*-based RNAi systems. Active transport of endogenous/viral proteins likely involves sophisticated interactions at the cytosolic side of the desmotubule, a compressed cortical ER segment linking adjacent cells through the PD lumen (Beachy and Heinlein, 2000; Kim et al., 2005). Strikingly, PD-based free GFP movement is desmotubule-independent (Oparka et al., 1999). Other, perhaps unique sRNA-intrinsic features could underpin their crossing of various PD-based boundaries as part of a potentially dedicated transport mechanism, including their negative charge if one assumes that si/miRNAs move as protein-free entities.

### Genetically tractable, discrete loci enable studies of endo-siRNA movement and functions in intact plants and perhaps over generations

Long-distance endo-RNAi studies involving grafted *dcl234* recipient tissues concluded that only a third of all *Arabidopsis* 24-nt siRNA-generating loci apparently spawn mobile species (Lewsey et al., 2016; Molnar et al., 2010), providing further possible evidence of selective siRNA mobility (Maizel et al., 2020). Recent work argues against this notion in cell-to-cell movement settings, however (Devers et al., 2020), and we further suggest here that all siRNAs have a graft-transmission potential. Yet, the dramatic albeit artefactual loss-of-AGO4 stability suggests that, for them to be detected in *dcl234*-receiving-rootstocks, siRNAs should be sufficiently abundant to stabilize their effectors and, reciprocally, for those effectors to stabilize their cargoes. Consequently, the amount, diversity and breath-of-activities of mobile endo-siRNAs have been likely underestimated in siRNA-deficient *dcl234* recipient tissues. Use of this now-apparent suboptimal background has been mandated by the domination, in movement studies, of TE/repeat-derived siRNAs (Lewsey et al., 2016; Molnar et al., 2010) whose genome-multiplicity makes single-loci knockouts *de facto* unavailable.

*IR71* illustrates how discrete, genetically-tractable sources of mobile endo-RNAi bestow the use of WT recipient-tissues to overcome *dcl234-*associated caveats, and how stress-induced endo-RNAi modulation can bypass unidirectionality caused by micro-grafting. The naturally vasculature-restricted *IR71* expression allowed mobile endo-RNAi study in intact plants by avoiding the contentious process of (micro)grafting altogether (Kehr, 2013). Mechanical tissue-separation uncovered a cell-to-cell, in addition to long-distance, component to *IR71-*siRNA mobility, revealing transgenic RNAi-like properties (Fig. 5; Devers et al., 2020)*. IR71*-siRNA cell-to-cell spread elicits RdDM, at least *in cis*, and *trans*-regulates dozens of mRNAs collectively promoting SAR *via* salicylate biosynthesis/signalling/metabolism (Pieterse et al., 2009). Conversely, downregulated (in all likelihood indirectly) mRNAs in *ir71* plants are biased toward the SA-antagonistic JA pathway (Pieterse et al., 2009). Its basal expression and non-autonomous action in healthy plants thus suggest that *IR71* could potentially act as a rheostat enabling rapid, reversible tuning of JA *vs* SA responses not achievable by separate transcriptional control of possibly dozens of discrete loci. The HS induction of *IR71* might also illustrate a trade-off for resource allocation between abiotic *vs* biotic stress adaptation (Berens et al., 2019).

We failed to identify the most upstream regulatory nodes – and thus the likely direct targets – of the *IR71*-controlled gene network. Indeed, transcriptome mining coupled to target predictions were not backed-up by degradome-based searches for AGO:sRNA-mediated slicing events. We note the unusual preponderance of 22-nt-long, instead of 21-nt-long, PTGS-inducing *IR71*-siRNAs. Recently, stress-induced 22-nt siRNAs were found to mostly incur translational repression of predicted mRNA targets in *Arabidopsis* (Wu et al., 2020). A similar effect from *IR71*-siRNAs would not have been diagnosed by transcriptomics reporting, perhaps, mere secondary consequences of dysregulating, at the translation level, a narrower subset of primary *IR71* targets. Regardless, and while more work is needed to assess *IR71’s* potential roles in SAR/stress regulation, our findings now provide a tangible framework for functional mobile endo-RNAi studies in *Arabidopsis*. The HS-induction of *IR71-* siRNA and their ensuing shootward movement in *ir71*/WT micro-grafting might also facilitate transgenerational epigenetic inheritance studies from this genetically-tractable locus.

## ACKNOWLEDGMENTS

We thank Voinnet lab members for fruitful discussions, A. Imboden for plant care, E Truernit for providing *pSUC2::tmGFP9* and *pSUC2::GFP* seeds, C. Himber for selecting homozygous *pATML1::SUL-IR* lines, S. Mirlohi and G. Schott for some images and the TRV-PDS infections, and the ETH ScopeM unit for providing the microscopy facility. This work was supported by an EMBO Long-Term Fellowship (ALTF 728-2009) to C.A.B., a Marie Curie Intra-European Fellowship for career development (FP7-PEOPLE-IEF; number 623826) to E.A.D. and a European Research Council advanced grant (Frontiers of RNAi-II; number 323071) to O.V.

## AUTHORS CONTRIBUTIONS

C.A.B., E.A.D., and O.V. designed the project and all experiments. C.A.B. conducted all experiments relating to *SS* grafting, MeSelect and *IR71*. E.A.D. conducted experiments relating to PD and *iCals3m*-induced callose deposition and *AS* grafting. A.S. performed all bioinformatic analysis. P.L. assisted in molecular analysis of *IR71*. S.L. and Y.H contributed the original pSUC2-based *iCals3m* system. C.A.B., E.A.D. and O.V. analyzed the data. C.A.B. and E.A.D. assembled all of the figures and, together with O.V., wrote the manuscript. All authors read and approved the manuscript.

## Material and Methods

### Plant material, growth conditions

Wild-type(WT) and mutant Arabidopsis (ecotype Columbia: Col-0) were germinated and grown under long-day light conditions at 22°C on ½ Murashige and Skoog(MS) medium. The *pSUC2::SUL-IR*(*SS*), *pIR71::GUS*, *pSUC2::GFP*, *pSUC2::tmGFP9, JAP3* and *dcl2-1/dcl3-1/dcl4-2* lines were described previously (Deleris et al., 2006; Devers et al., 2020; Himber et al., 2003; Smith et al., 2007; Stadler et al., 2005). The T-DNA-insertion line for *IR71* is SALK_006309. For root isolation, plants were either grown vertically on mesh-covered solid media (Sefear Nitrex 03-65125) or hydroponically (Devers et al., 2020; Gibeaut et al., 1997; Tocquin et al., 2003). For iCals3m experiments the *pSUC2::iCals3m*(*iSC*) parental lines “21-1”, “23-1” and “25-1” were tested for callose induction and developmental defects on a 0-100 µM 17-ß-estradiol (Sigma) concentration-range. Since “23-1” and “25-1” showed growth defects at all concentrations, “21-1”(*iSC*) was selected and crossed to *SS*. Seeds of *iSC* or *iSCxSS* (F3) were germinated and upon ermegence of the first true leaves L1&L2, ∼10-15 seedlings were transferred to fresh solid MS containing 20µM 17-ß- estradiol or DMSO (Sigma). For heat-shock experiments, ∼8 week-old micro-grafted plants were subjected to 37°C for 20hrs.

### Micrografting

Micro-grafting was as previously described (Brosnan et al., 2007; Turnbull et al., 2002). Seedlings were germinated on vertical ½ MS medium plates for 4-6 days. Transverse cuts were made within a millimeter to the shoot apex and subsequently aligned on a 0.45µm nitrocellulose filter (Millipore) on top of two moist Whatman no.1filter paper’s layers. Sealed plates were positioned vertically for 7 days prior to transfer to hydroponic or normal ½ MS media.

### Cloning procedures

Cloning *pSUC2::iCals3m* involved multi-site gateway (Invitrogen). Entry vector *p1R4-pAtSUC2::XVE* (Siligato et al., 2016), *pDONR221-cals3m* (Vatén et al., 2011) and *p2R3a-3AT/nosT* (Siligato et al., 2016) were recombined with the destination binary vector *pCAM-hyg-R4R3* (Siligato et al., 2016). The CC-specific promoter *pSUC2* was PCR-amplified from genomic DNA (primers in table S4) and recombined into the Gateway vector *pDONR P4-P1R*. For cloning *pATML1::SUL-IR,* the ATML1 promoter was PCR-amplified (primers ATML1promoter-F and ATML1promoter-R;Table S4) and inserted as an EcoRI-XhoI restriction fragment into *pFGC5941.* All final constructs were introduced into *A. tumefaciens* GV3101 for *Arabidopsis* transformation via floral dip (Clough and Bent, 1998).

### Immunoprecipitation

Root tissue was ground in IP buffer (50mM Tris-HCL, pH7.5, 150mM NaCL, 10% glycerol, 0.1% NP40) containing cOmplete protease inhibitor (Roche). After clearing lysates at 12,000x g (10 min), supernatants were transferred into fresh tubes and 10% vol used as input. Lysates were incubated with anti-AGO4 antibody (1:10000; Agrisera AS09 617) for 1-2hrs, upon which protein A/G magnetic beads (Pierce) were added before further incubatation for 1-2hrs. Beads were washed 3x with IP buffer, proteins acetone-precipitated from the organic phase and RNA extracted as described below.

### Protein extraction, gel blot analysis

Proteins were resolved via SDS-PAGE, transferred to Immobilon-P PVDF membranes (Millipore) and subjected to the anti-AGO4 antibody (1:5000; Agrisera AS09 617) in blocking buffer (1x PBS, 0.1% Tween-20, 5% skim-milk powder). After washing, membranes were incubated for 1hr at RT with a horseradish peroxidase-conjugated anti-rabbit goat serum (1:10000; Abcam ab6721). After washing, detection was through the ECL Western Blotting Detection Kit (GE Healthcare) revealed via ChemiDocTouch from Bio-Rad. Coomassie blue-stained membranes provided loading controls.

### RNA isolation, cDNA synthesis and qRT-PCR

Tri-Reagent was used for total/immunoprecipitated RNA extractions. Reverse transcription of DNaseI-treated RNA was performed with the Maxima First Strand cDNA Synthesis Kit (Thermo). qRT-PCR was performed on LightCycler480II (Roche) using the KAPA-SYBR-FAST qPCR kit (Roche; Brosnan et al., 2019). Ct-values were determined by 2^nd^-derivative max on average of technical triplicates. Data are presented as average of three biological replicates calculated from ΔCt values obtained by comparing the gene-of-interest to a set of house-keeping genes. Normal distributions were confirmed by Shapiro-Wilk test (α =0.02). Equal variances were evaluated by F-test (95% confidence interval).

### RNA gel blot analysis

Low-molecular-weight northern blots were conducted as described (Brosnan et al,. 2019). Total/immunoprecipitated RNA was resolved on 17.5% denaturing PAGE, electrotransferred to HyBond-Nylon NX (GE Healthcare) and EDC-crosslinked (Pall and Hamilton, 2008). Probes against individual miRNAs were generated by end-labelling with [γ-^32^P]-dATP of complementary oligonucleotides. Probes against siRNA populations were PCR products labelled with the Prime-a-Gene kit (Promega). Over-night hybridization at 42°C was in PerfectHyb buffer (Sigma) followed by washing four times with 2x SSC 0.1% SDS at 50°C. Signals were revealed with a Typhoon FLA9500 scanner (GE Healthcare). For re-probing, membranes were stripped in boiling 0.1% SDS.

### Bisulfite sequencing

As described (Pumplin et al., 2016), total genomic DNA was converted using the EZ DNA Methylation Kit (ZYMO Research) and amplified with primers listed in Table S4. The amplified DNA was mobilized into pGEM-Teasy, with at least 10 independent clones sequenced per treatment. Analysis was performed using Kismeth (Gruntman et al., 2008) (http://katahdin.mssm.edu/kismeth/revpage.pl). 95% confidence intervals were calculated using Wilson score (Table S1).

### Small RNA sequencing

RNA from *SS*-WT micro-grafting was processed into libraries using modified Illumina protocols by -and sequenced at-Fasteris (http://www.fasteris.com, Switzerland) using the HiSeq2000 sequencer. For WT and *ir71* mutant libraries, RNA (10µl at 200ng/µl) was processed into libraries and sequenced on the HiSeq2500 sequencer at the Functional-Genomics-Center-Zurich (http://www.fgcz.ch/).

### mRNA sequencing

Total RNA from WT and *ir71* epidermis was sequenced (random hexamer paired-end 2x125-bp-stranded RNA-sequencing) at Fasteris (http://www.fasteris.com, Switzerland) after ribosomal depletion.

### sRNA sequencing data processing and representation

Small RNA data from grafting experiment are from GEO accession GSE112885 and have been processed as described in Devers et al. 2020. For WT and *ir71* mutant libraries, reads were clipped using fastx_clipper from fastx-toolkit (http://hannonlab.cshl.edu/fastx_toolkit/) looking for the adapter sequence’s first 15- nt(TGGAATTCTCGGGTG). 17-to-30-nt reads with identical sequences were grouped using the `processReads` function from the ncPRO-seq pipeline (Chen et al., 2012) and aligned against the *A. thaliana* genome (TAIR10) using Bowtie2 (Langmead and Salzberg, 2012) with options `-k 100 --score-min L,0,0`. In Fig. S6, simple genomic position comparison was applied to retrieve sRNA read counts and positions corresponding to *SS* or *IR71*, subsequently used to calculate normalized read-counts (reads per 1millions mapped reads) for each nucleotide and to produce a graphical representation with R. Color-coded reads-lengths are indicated in figures’ legends.

### Quantification of TE-derived, graft-transmitted sRNAs

To quantify TE-derived sRNA graft-transmission, we used data from Molnar et al., 2010 (GSM518438-40, GSM518443-45). Raw data were retrieved from GEO/SRA, adaptor sequences removed using fastx_clipper (-a TCGTATGCCGT) and processed as described above. TE being repeated sequences, in order to quantify the sRNA responsible for the signal obtained in Fig. 4A we first aligned the sequences corresponding to the probes to the TAIR10 genome using bowtie2. Then, sRNA reads overlapping the inverse strand of the obtained region, enlarged by 5bp on both sides, were selected using intersectBed from bedtools (v2.25.0; Quinlan and Hall, 2010) with options `-S -F 0.9`. For each region, read counts were normalized (reads per million mapped reads) and the average between replicates was represented as boxplot with R. Mean values for each type of TE sequences were indicated in red.

### mRNA sequencing data processing

Reads were trimmed using Trimmomatic (v0.36; (Bolger et al., 2014)) with parameters `ILLUMINACLIP:TruSeq2-PE.fa:2:30:10 SLIDINGWINDOW:4:15 MINLEN:50` and mapped to TAIR10 Ensembl-genome-and-genes-annotation (http://ccb.jhu.edu/software/tophat/igenomes.shtml) using TopHat (v2.0.11; (Trapnell et al., 2009) and Bowtie2 (v2.2.1.0; Langmead and Salzberg, 2012). Counts for annotated genes (Ensembl TAIR10) were generated using the `featureCounts` function from Subread (Liao et al., 2014) with options `-S rf -p -s 1 -C -B -O`. Differential analysis and MAplot in Fig. 5D were done with DEseq2 (v1.12.4; (Love et al., 2014) taking into account the two replicates. Genes with adjusted p-values <0.01 were considered differentially expressed.

### Gene Ontology enrichment

Up- and down-regulated genes were subjected to agriGO gene-ontology-enrichment analysis (Tian et al., 2017) using the TAIR10 *Arabidopsis* reference genome and default parameters. Enriched GO terms were then transferred to REVIGO (Supek et al., 2011) with parameters: `Similarity Medium ; database : Arabidopsis thaliana ; semantic similarity measure : Resnik (normalized)`. Only terms with a frequency <1 were considered. Histograms in Fig. 5E-F represent the proportion of genes in the query list and background for the top 10 ontology terms following ranking based on log_10_ (p-value).

### Leaf epidermis peeling (Meselect)

As described (Svozil et al., 2016), the lower epidermis was removed by peeling after placing leaves between two sticky tapes. Residual mesophyll cells were removed in protoplasting solution (1% cellulase Onozuka RS (Yakult), 0.4M mannitol, 10mM CaCl2, 20mM KCl, 0.1% BSA and 20mM MES, pH 5.7). The epidermal side was incubated for 15min and the vasculature for 20-30min; separated tissues were washed twice with ice-cold wash buffer (154mM NaCl, 125mM CaCl2, 5mM KCl, 5mM glucose and 2mM MES, pH 5.7). The vasculature was removed with forceps and frozen in liquid nitrogen. RNA was extracted as described above.

### Callose staining

Plants were fixed in acetic acid/ethanol 1:3 (v/v) for at least 2hrs and washed three times for 20min in 150mM K_2_HPO_4_. Aniline blue 0.01% (w/v) in 150mM K_2_HPO_4_ was used to stain callose depositions overnight at RT protected from light. Stained callose structures were imaged using an epifluorescence microscope with a DAPI filter.

### Data Availability

Data have been deposited into the Gene Expression Omnibus (GEO) under the accession number GSE112929 for the RNA-sequencing data from WT and ir71 epidermis and under the accession number GSE112861 for the sRNA-sequencing from WT and ir71 shoot apex.

### Supplementary Figure Legends

**Figure S1. Photobleaching phenotype in source versus sink leaves of *SS* transgenic Arabidopsis.** 4 weeks old soil grown *Arabidopsis* plant with typical photobleaching phenotype due to RNAi triggered by expression of *pSUC2::SUL-IR* (*SS*). The letters indicate the position of the extracted leaves on the intact plant. From oldest to youngest leaf A corresponds to L5 and B to L11 and were designated as source leaf (SoL) and sink leaf (SiL), respectively. Roman numerals and black arrows indicate the leaf veins of class I (mid-rib), II (veins of second order, originating from the mid-rib) and III (veins of third order, originating from class II veins). Scale bars represent 5 mm. For better comparison, the background in those pictures was masked but original photos used in this figure can be found as supplementary data files.

**Figure S2. Phenotypic quantification and distribution of callose depositions at vascular bundles in *iSC* expressing lines**. **(a)** Phenotypic scoring of the *SS* bleaching phenotype of each leaf stage in populations of *iSC* (left) and *SS* (right) plants. Yellow bars represent the percentage of the population showing vein centered bleaching of chlorophyll, green bars represent the percentage of the population showing no signs of vein centered bleaching and grey bars represent the percentage of the population having the respective leaf missing at the time of screening. Error bars represent SE, n = 4 (*iSC*) and 7 (*SS*) populations of 6-12 plants. **(b)** Epifluorescence images of an aniline blue stained leaf series of *iSC* plant lines grown on either control (left) or inducing (right) media. Concentrations correspond to 17-ß-estradiol present in the media. Scale bars represent 200µm. Co = cotyledon; L = leaf. **(c)** qRT-PCR analysis of endogenous *SUC2* and *DCL4* mRNA levels in mock- or ß-estradiol-induced seedlings. Error bars represent SE, n = 6-10. Asterisks indicate statistically significant differences (p≤ 0.05, two-tailed Student’s t-test with equal variance [F-test, p > 0.05]).

**Figure S3. Representative images equivalent aged flowering WT (left), *SS* (middle) and *amiRSUL* (right) plants.** Insets show lack of photobleaching in stem tissue of each respective genotype.

**Figure S4. Comparison of Argonaute transcript levels in various tissues and cell types of *Arabidopsis*.** Analysis of publicly available transcriptome data to compare transcript levels of AGO1, 2, 4 and 6 in **(a)** different plant tissues using the EMBL-EBI Gene expression atlas (https://www.ebi.ac.uk/gxa/home; release 37) (source data from PMID: 23136377) or **(b)** in different leaf cell-types manually extracted from data published in Berkowitz et al., 2021.

**Figure S5. Analysis of sequenced graft-transmissible sRNA populations.** Boxplot representation of sRNA abundance (read per millions mapped reads) corresponding to 3 classes of repeat sequences (Rep2, 1003 and SimpleHat). Sequencing libraries correspond to graft-transmissible sRNA from Molnal et al., 2021. WT/WT refers to Col/Col and dcl234 refers to Col-dcl2,3,4 of the original nomenclature.

**Figure S6. siRNA profiles of mobile *SS* populations and *IR71* knockout loci. (a)** Profile and histograms of *SS*-derived siRNA reads from grafted WT scion (Top) on top of a *SS* rootstock (Bottom). **(b)** Upper panels show the profile and size distribution (right) of IR71-derived siRNAs in WT plants. Middle panel shows siRNA profile of IR71-derived siRNAs in T-DNA line (*ir71 -/-*). The lower panel shows the siRNA profile of IR71-derived siRNAs in T-DNA line (*ir71 -/-*) with adjusted scale demonstrating background levels of siRNA. Top and middle panels are also shown in Figure 4. RPM – Reads per Million, color code: Blue – 20-21-nt, Green – 22-23-nt and Red – 24-25-nt.

