## Supplementary data for "*In planta* dynamics, transport-biases and endogenous functions of mobile siRNAs in *Arabidopsis*"

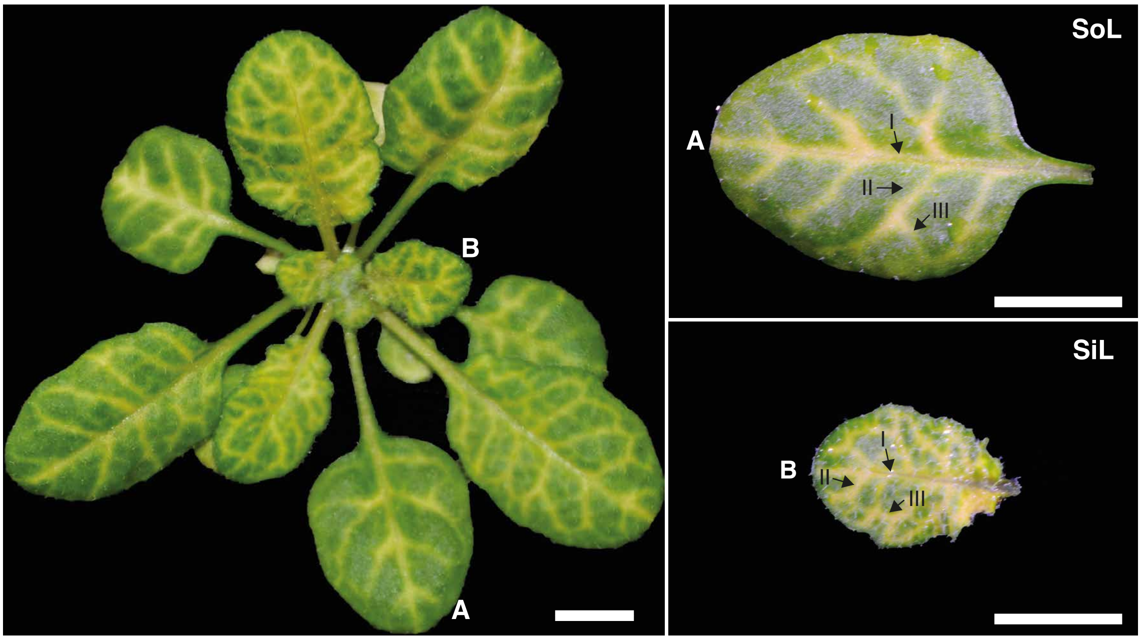


**Supplementary figure 1**

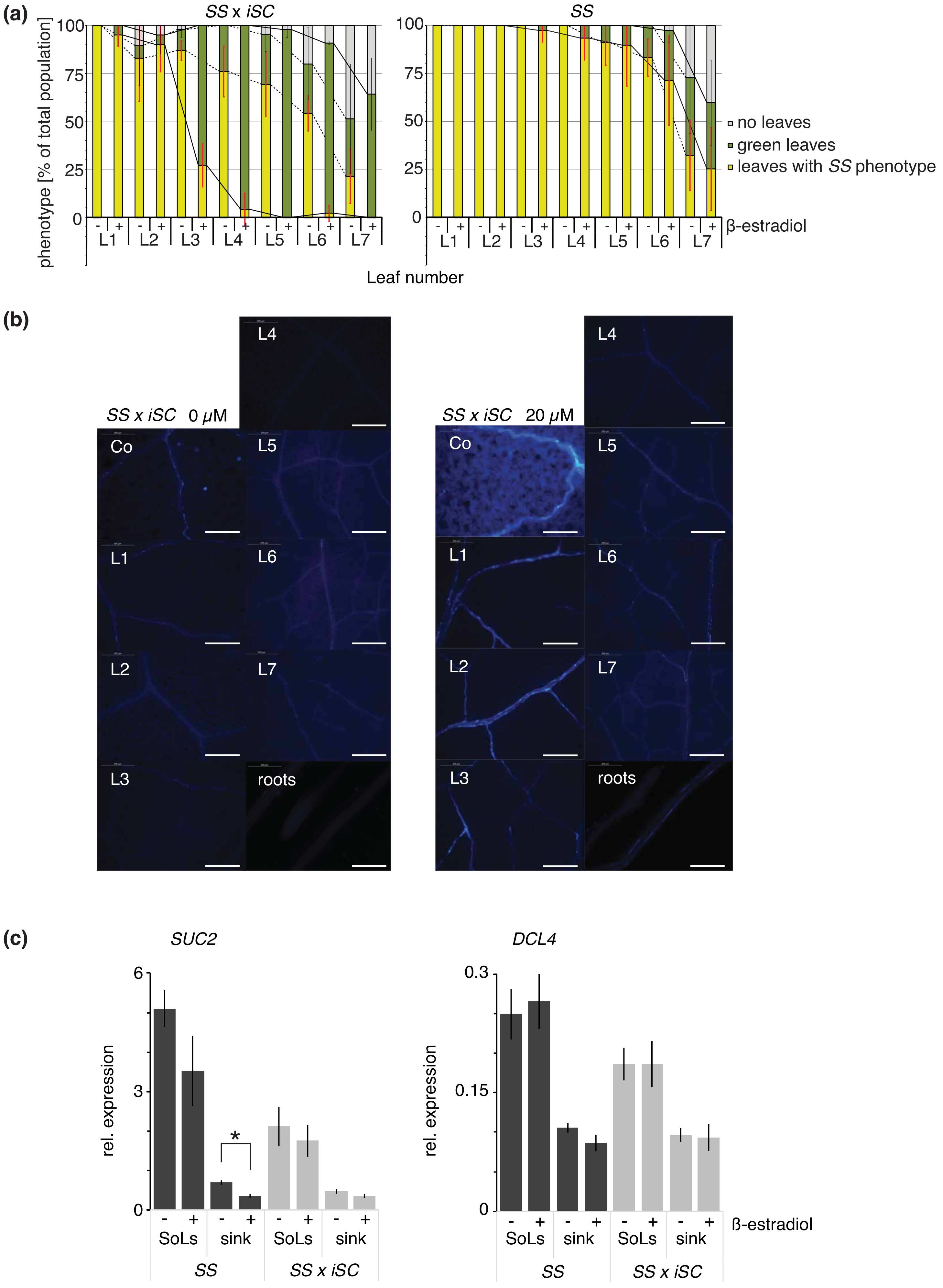


**Supplementary figure 2**

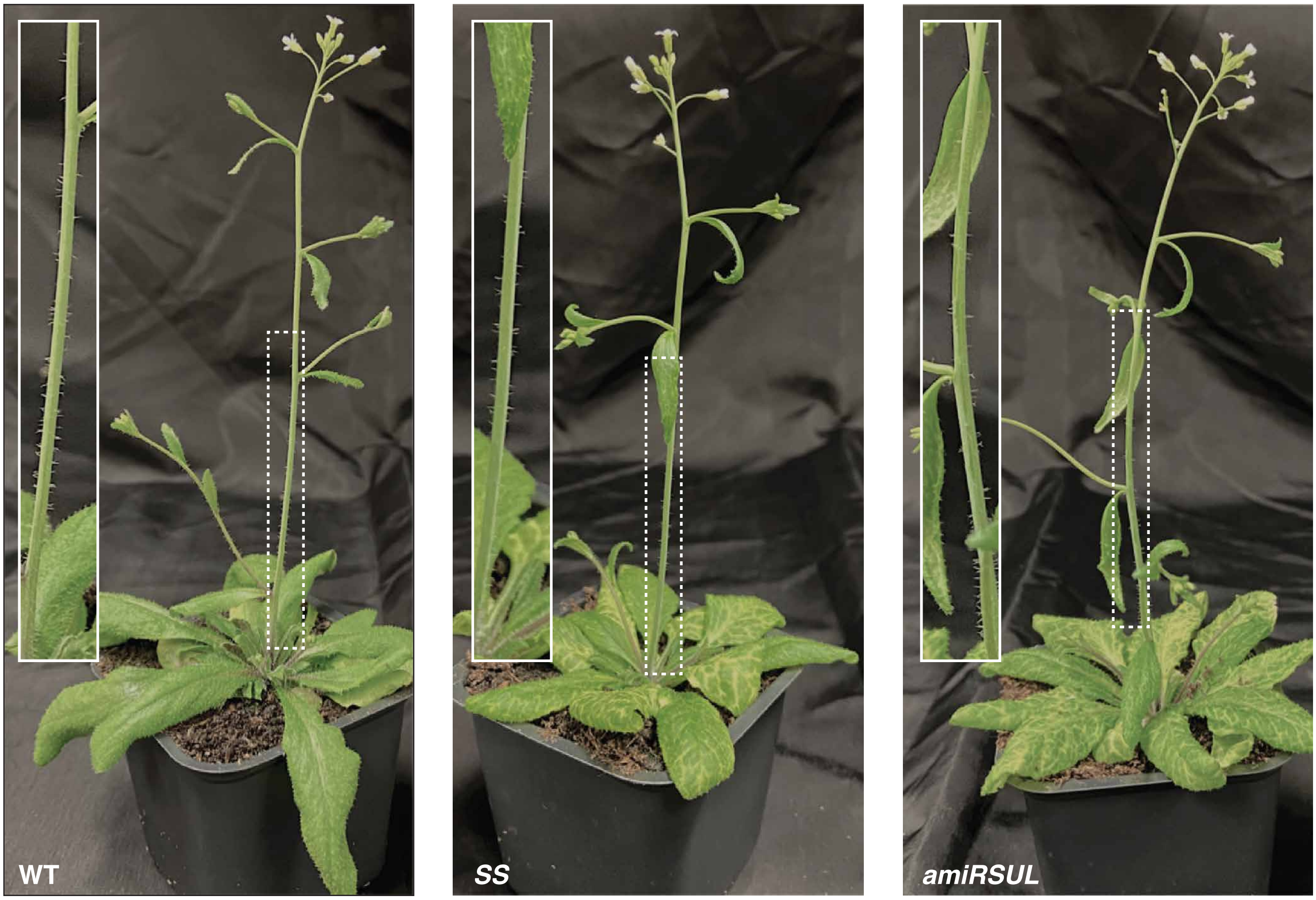


**Supplementary figure 3**

**(a)**

Expression level in TPM

0 – 197

| | leaf | flower | fruit | root |
|---|---|---|---|---|
| AGO1 | 112.0 | 197.0 | 125.0 | 150.0 |
| AGO4 | 31.0 | 99.0 | 67.0 | 32.0 |
| AGO2 | 8.0 | 16.0 | 22.0 | 12.0 |
| AGO6 | 2.0 | 4.0 | 4.0 | 3.0 |

**(b)**

Expression level in TPM

0 – 50.15

| | whole leaf | epidermis | mesophyll | vasculature |
|---|---|---|---|---|
| AGO1 | 50.15 | 11.57 | 7.16 | 24.14 |
| AGO4 | 22.98 | 6.67 | 1.37 | 16.89 |
| AGO2 | 5.72 | 1.99 | 0.51 | 4.02 |
| AGO6 | 1.15 | 0.38 | 0 | 0.87 |

**Supplementary figure 4**

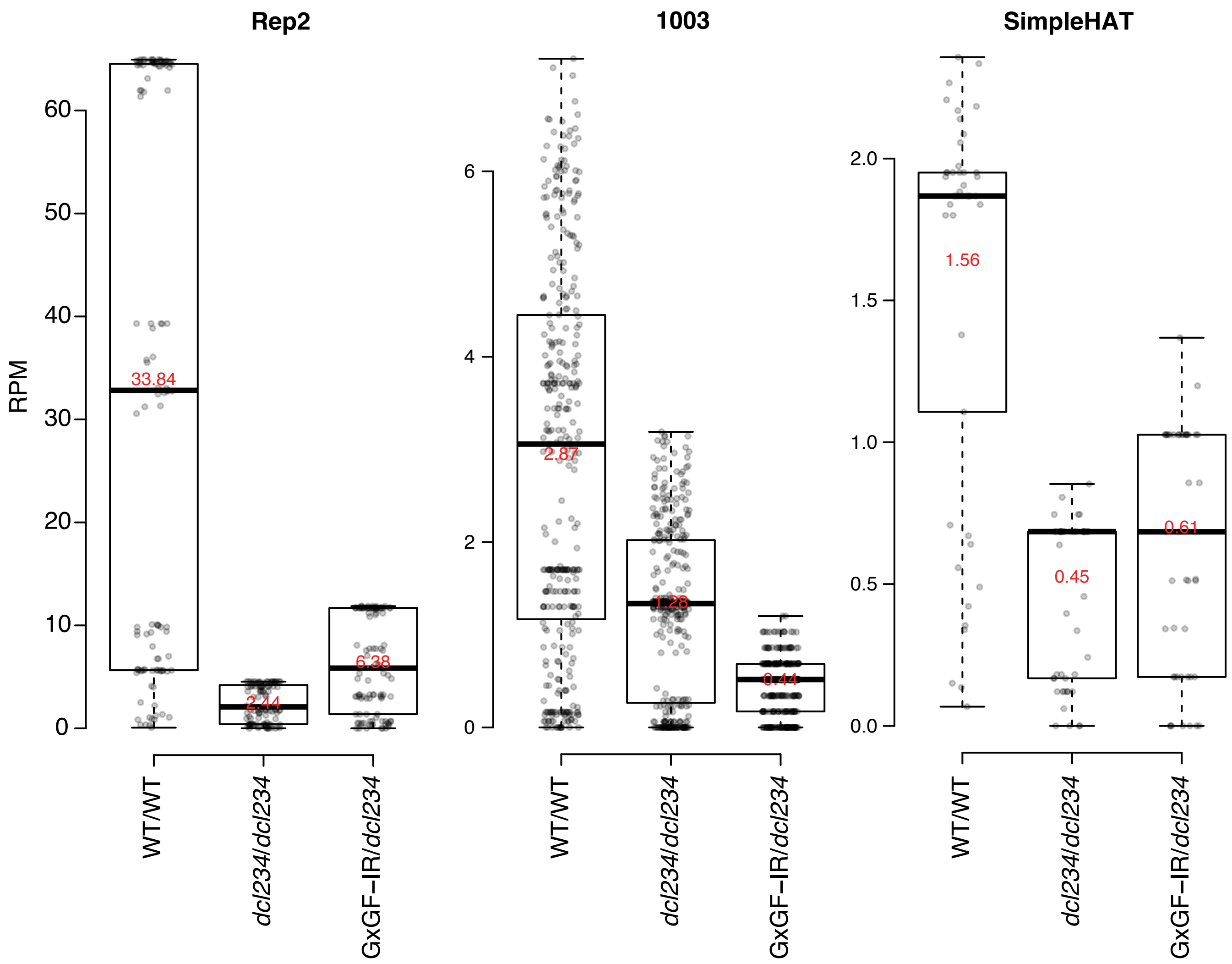


**Supplementary figure 5**

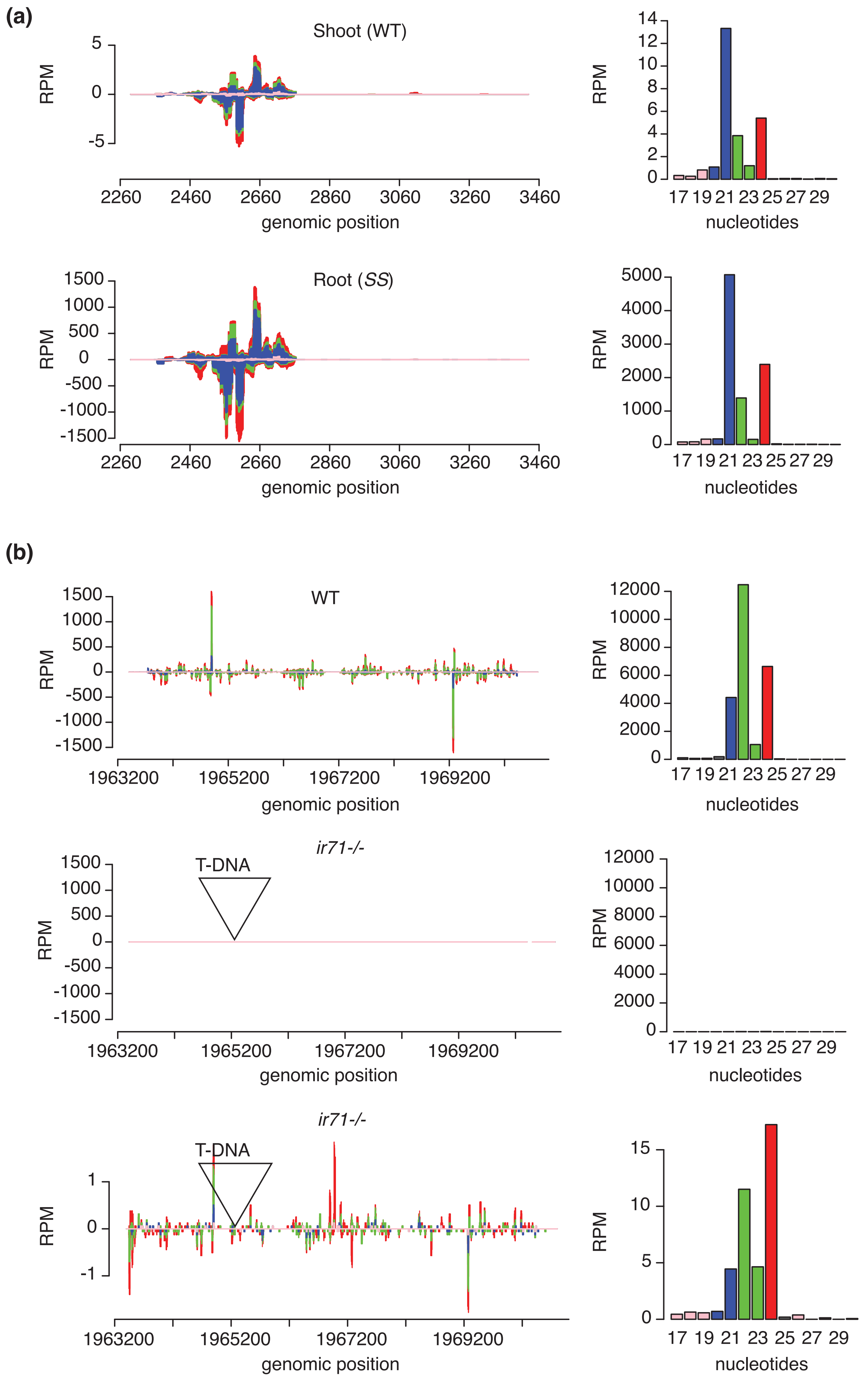


**Supplementary figure 6**
